# An HSP90-regulated *reduced-eye* phenotype in *Tribolium* shows fitness benefits and thus provides evidence for evolutionary capacitance

**DOI:** 10.1101/690727

**Authors:** Rasha Aboelsoud, Joachim Kurtz

## Abstract

Evolution relies on genetic variation as the raw material for adaptation. The release of cryptic genetic variation (CGV), which can be regulated by the evolutionary capacitor heat shock protein 90 (HSP90), may thus be important for rapid adaptation. However, the fitness benefits of HSP90-regulated phenotypes are still under debate. Here, we show in the important model insect *Tribolium castaneum* that HSP90 impairment by two independent methods, RNA interference and chemical inhibition, revealed the same *reduced-eye* phenotype, which was stably inherited without further HSP90 inhibition. The penetrance and fitness of this trait increased under ambient light stress. This is the first demonstration that a phenotype released through HSP90 inhibition can be adaptive.

## Introduction

Canalization was described as the state of a phenotype’s stability against genetic and environmental perturbations (Lack et al., 2016; Waddington, 1942). Heat shock protein 90 (HSP90) facilitates canalization, as it buffers against morphological variation by biochemical, but maybe also by further genetic and epigenetic mechanisms (Rutherford and Lindquist, 1998; Sollars et al., 2003). HSP90 is primarily a molecular chaperone that maintains protein homeostasis by assisting correct protein folding (Lindquist and Craig, 1988; Taipale et al., 2010). Many client proteins of HSP90 are involved in the regulation of development (Rutherford et al., 2007). Consequently, impaired HSP90 function, for example in *Drosophila Hsp83* mutants, results in a multitude of morphological alterations (Rutherford and Lindquist, 1998). As artificial selection has been shown to maintain these altered phenotypes in subsequent generations, HSP90 has been described as an evolutionary capacitor, referring to its ability to store and release genetic variation (Queitsch et al., 2002; Rutherford and Lindquist, 1998; Sangster et al., 2008b). As long as HSP90 prevents that certain genetic variants are expressed phenotypically (i.e., maintains genetic variation in a cryptic state), selection will not act on such variants and phenotypic evolution will be temporarily constrained (Flatt, 2005; Rohner et al., 2013; Sangster et al., 2008a). However, the availability of HSP90 may become limited, for example under stressful environmental situations, when HSP90 is needed as a chaperone by many proteins (Borkovich et al., 1989; Chen and Wagner, 2012; Peuss et al., 2015). Under such conditions, stored genetic variation is released, expressed as phenotypic differences, and selection can act on it. Therefore, HSP90 could be a molecular mechanism underpinning the de-canalization and potential subsequent assimilation of traits (Queitsch et al., 2002; Rutherford and Lindquist, 1998).

Additional functions of HSP90 have been described that might likewise support its role in facilitating rapid adaptation. HSP90 also functions as a regulator of gene expression via epigenetic mechanisms, chromatin remodeling (Sollars et al., 2003; Tariq et al., 2009), and the regulatory effects of endogenous retroviruses on nearby genes (Hummel et al., 2017). HSP90 reduction has also been suggested to lead to transposon activation, thereby generating new genetic variation (Specchia et al., 2010). Only very few studies have so far addressed the potential adaptive value of HSP90-regulated phenotypes (Chen et al., 2012; Koubkova-Yu et al., 2018; Queitsch et al., 2002; Rohner et al., 2013). For example, in surface populations of cavefish *Astyanax mexicanus*, HSP90 inhibition induced variation in eye size, suggesting that the evolution of adaptive eye loss in the dark might have been facilitated by HSP90 in nature (Rohner et al., 2013). Only one study directly focused on fitness, concluding that the correlative fitness costs of lines selected for an HSP90-released *deformed eye trait (dfe)* in *D. melanogaster* were low (Carey et al., 2006), while the adaptive benefits of the trait itself under any imaginable environment were considered unlikely. In the red flour beetle, *Tribolium castaneum* we recently found consistent down-regulations of *Hsp83* and *Hsp90*, two HSP90-coding genes, mediated by social cues mimicking a stressful environment (Peuss et al., 2015). This implies that the level of HSP90 might be adaptively regulated according to whether or not the release of cryptic genetic variability may be advantageous. We therefore hypothesized that HSP90 reduction in this important model organism releases selectable cryptic morphological variants that might have a fitness benefit under specific environmental conditions. We thus suppressed HSP90 function in *T. castaneum* via RNAi and chemical inhibition and examined phenotypic variation in the offspring, as well as trait inheritance and fitness effects in subsequent generations.

## Results

### HSP90 inhibition releases a reduced-eye phenotype

We determined the effect of *Hsp83-*reduction in a genetically variable wild-type strain of *T. castaneum* (Cro1), using RNA interference (RNAi). *Hsp83* knockdown, confirmed by RT-qPCR, resulted in a strong, dose-dependent reduction in the injected males and females, while there were normal *Hsp83* levels in their offspring, indicating the depletion of the introduced dsRNA (Figure 1-figure supplement 1A and B). The maternal treatment with the low dsRNA concentration reduced the offspring number, while the high dsRNA concentration completely sterilized the females due to its effects on ovarian maturation and oocyte development, as previously shown (Xu et al., 2010, 2009).

As we were interested in potential evolutionarily relevant effects of parental HSP90-reduction on offspring, we examined their phenotypes. We observed morphological variations (Figure 1-table supplement 1) such as abnormal larval eyes (ocelli), missing leg segments, variant body pigmentation, and larval malformations (Figure 1A and B). While these larval abnormalities were not maintained into the adult stage (Figure 1-figure supplement 2A), leg malformations occurred in up to 4% of the adults (Figure 1-figure supplement 2B and C).

**Figure 1.**
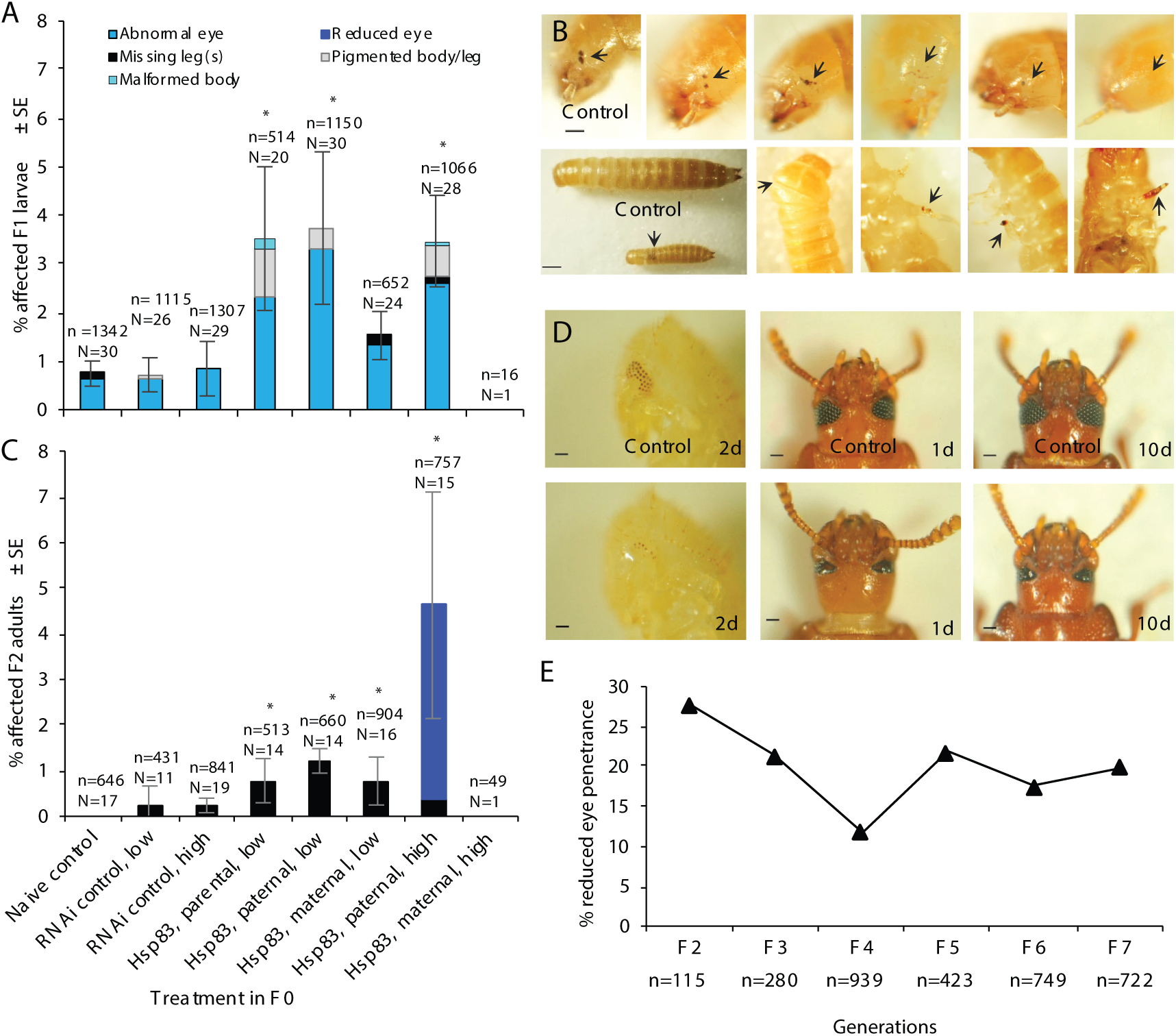
Morphological variations in F1 and F2 offspring of HSP90-inhibited parents (F0); RNAi-mediated knockdown of *Hsp83* with 20 (low) and 100 (high) ng/μl dsRNA. (**A**) F1, proportion of affected larvae (total number of *n* screened individuals per treatment produced from *N* pairs shown above each bar; significant differences indicated by asterisks, *p*<0.05, relative to naïve control; see details in Figure 1-table supplement 1). (**B**) F1 larvae, example phenotypes indicated by arrows: abnormalities in larval eyes (distant, scattered, reduced or absent ocular lobes), and other larval body parts (pigmentation, malformed or missing appendages); scale bars for the eye: 100 µm and for whole larva:1000 µm. (**C**) F2, proportion of affected adults produced from *N* pairs; *n =* total number of screened individuals. (**D**) F2 adults, reduced-eye phenotype: the development of compound eyes and progressive retina differentiation in control and affected (reduced-eye) beetles at different ages (2-day-old pupae, 1- and 10-day-old adults); scale bars: 100 µm. (**e**) Penetrance of the reduced-eye phenotype over successive generations; total number of *n* screened individuals per generation is shown under the X axis. *Hsp83* knockdown confirmation by RT-qPCR is shown in Figure 1-figure supplement 1. The morphological variations in F1 adults after *Hsp83* knockdown are shown in Figure 1-figure supplement 2.

When we selected and crossed (as single pairs) these F1 adults with a leg phenotype, we not only observed leg phenotypes in the resulting F2 adults, but an even higher proportion (more than 4%) of another phenotype, characterized by a strongly reduced eye size, due to a reduced number of ommatidia (32 of 757 scored F2 beetles; Figure 1C and D; Figure 1-table supplement 1). Affected individuals came from the same RNAi-treated grandfather and from two of the leg-affected families, in which about a quarter of individuals exhibited this reduced-eye trait: 25.5% (12/47) and 29.4% (20/68). To determine the persistence of this trait in the absence of selection, we continued breeding offspring from the affected families until the F7 generation and found a rather stable proportion (∼20%) of individuals with reduced eyes (Figure 1E).

To confirm that the reduced-eye phenotype was linked to the inhibition of HSP90, we used the HSP90-specific chemical inhibitor 17-DMAG (a water-soluble geldanamycin derivative) to inhibit HSP90 function (Jez et al., 2003). Larvae were placed on flour discs containing 10 (low) and 100 (high) µg/ml 17-DMAG and the inhibition of HSP90 function was confirmed by significant up-regulation of the stress gene *Hsp68a* (HSP70 protein family) (Kudryavtsev et al., 2017; Zhou et al., 2013) (Figure 2-figure supplement 1A). Two of 26 treated beetles showed reduced eyes and legs, but we could not obtain any offspring from these beetles. We thus paired 17-DMAG-treated, but morphologically normal beetles. We observed the reduced-eye phenotype described above in 0.4% (one of 226) and 5.1% (39 of 764) of their offspring, for the low and high parental 17-DMAG concentration, respectively (Figure 2-table supplement 1). The latter were derived from three of 13 families, with 22.2% (8/36), 22.0 % (9/41), and 16.6% (22/136) affected offspring. Reduced eyes were not found in either the negative control (i.e. flour discs without 17-DMAG) or the naïve control (i.e. feeding on normal flour) (Figure 2A).

**Figure 2.**
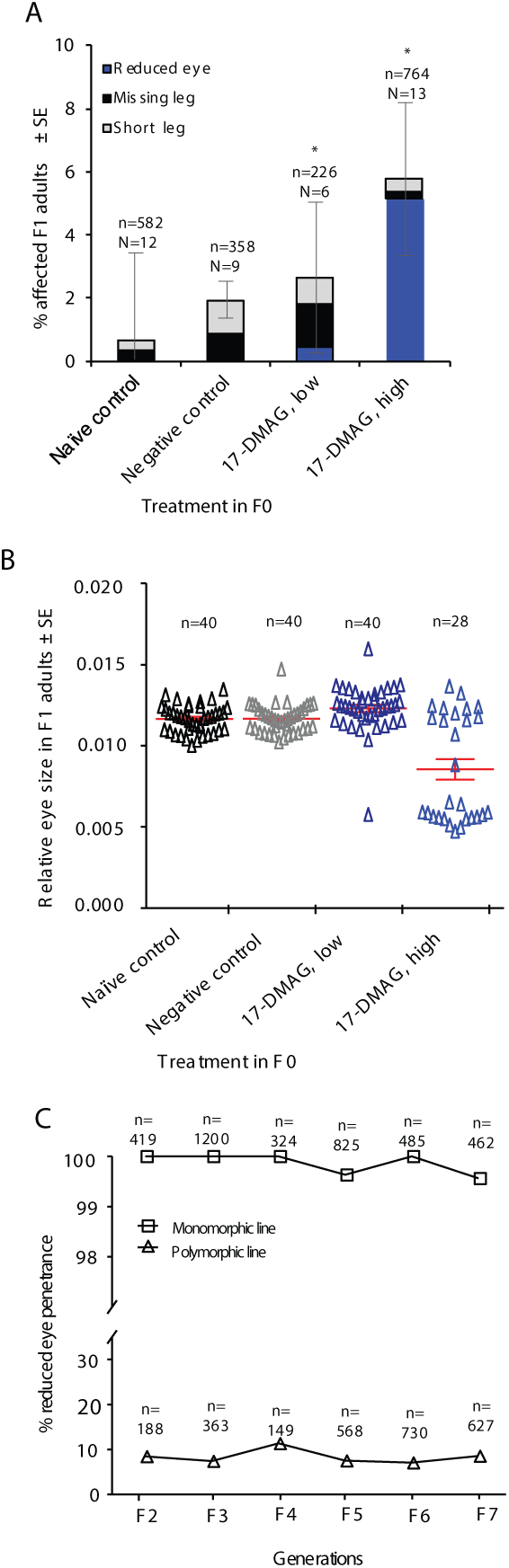
Morphological variations in F1 offspring of HSP90-inhibited parents (F0); chemical inhibition mediated by 17-DMAG at 10 (low) and 100 (high) μg/ml. (**A**) F1, proportion of affected adults (total number of *n* screened individuals produced from *N* pairs per treatment shown above each bar; significant differences indicated by asterisk, *p*<0.05, relative to negative control; see details in Figure 2-table supplement 1). (**B**) F1 adults, eye size relative to body length; each point shows the relative eye size of an individual with normal eyes (open dots) or reduced eyes (filled dots); mean ± SEM are shown in red. (**C**) Penetrance of the reduced-eye phenotype over seven generations in monomorphic and polymorphic lines; *n =* total number of screened individuals per generation; the monomorphic line was established by artificial selection for reduced-eye beetles and the polymorphic line by selection for normal-eye siblings. The artificial selection was carried out only in F1; no further selection was carried out in the following generations. The effect of 17-DMAG was tested via RT-qPCR of Hsp68 as a marker; *Hsp83* expression in beetles from the monomorphic lines was analysed by RT-qPCR; data are shown in Figure 2-figure supplement 1. The 17-DMAG experiment was repeated two times independently and the results are shown in Figure 2-figure supplement 2.

When we measured the eye size., we found that the mean eye sizes were 0.0289 mm^2^ ± 0.0006 in the high 17-DMAG concentration treatment, 0.0434 mm^2^ ± 0.0002 for the low 17-DMAG concentration, 0.0427 mm^2^ ± 0.0001 in the negative control and 0.0422 mm^2^ ± 0.0001 in the naïve control (Figure 2B). Beetles with reduced eyes did not differ in *Hsp83* expression from unaffected beetles (Figure 2-figure supplement 1B).

We then selected for the reduced-eye trait in this F1 generation and found almost 100% reduced eyes in all subsequent generations up to the F7 (Figure 2C). When we used unaffected siblings in the F1 for breeding, we consistently found less than about 10% of beetles with reduced eyes (Figure 2C) (these beetle lines will now be called *polymorphic*, to distinguish them from the *monomorphic* lines produced by selecting affected beetles in the F1). To further confirm our findings, we repeated the 17-DMAG treatment in an independent set of beetles and again found the reduced-eye phenotype in the F2 (Figure 2-figure supplement 2A). Moreover, to distinguish between maternal and paternal effects, in an additional experiment, we treated only females, only males, or both, and found beetles with reduced eyes in the F2 when both grandparents had been treated (Figure 2-figure supplement 2B).

### Inheritance of the reduced-eye phenotype

To examine whether the reduced-eye phenotype follows Mendelian inheritance, males were outcrossed with non-related, unaffected females. The trait disappeared in the F1 generation but was expressed again in 16-28% of the F2 offspring, which is consistent with the 25% expected for a recessive Mendelian trait (Figure 3A). The penetrance slightly differed in the F2 when affected females were outcrossed, ranging from 12-18% (after exclusion of two outliers), which might suggest additional, potentially sex-specific effects. We further asked whether the trait produced by *Hsp83* RNAi and the 17-DMAG treatment might be based on the same genetic locus. We thus used the monomorphic, reduced-eye lines produced by selecting affected offspring of beetles that had undergone these different treatments. We crossed affected individuals from the different lines and always observed 100% reduced-eye offspring in the F1, which is consistent with the expectation for one and the same recessive genetic locus being responsible in both lines (Figure 3B).

**Figure 3.**
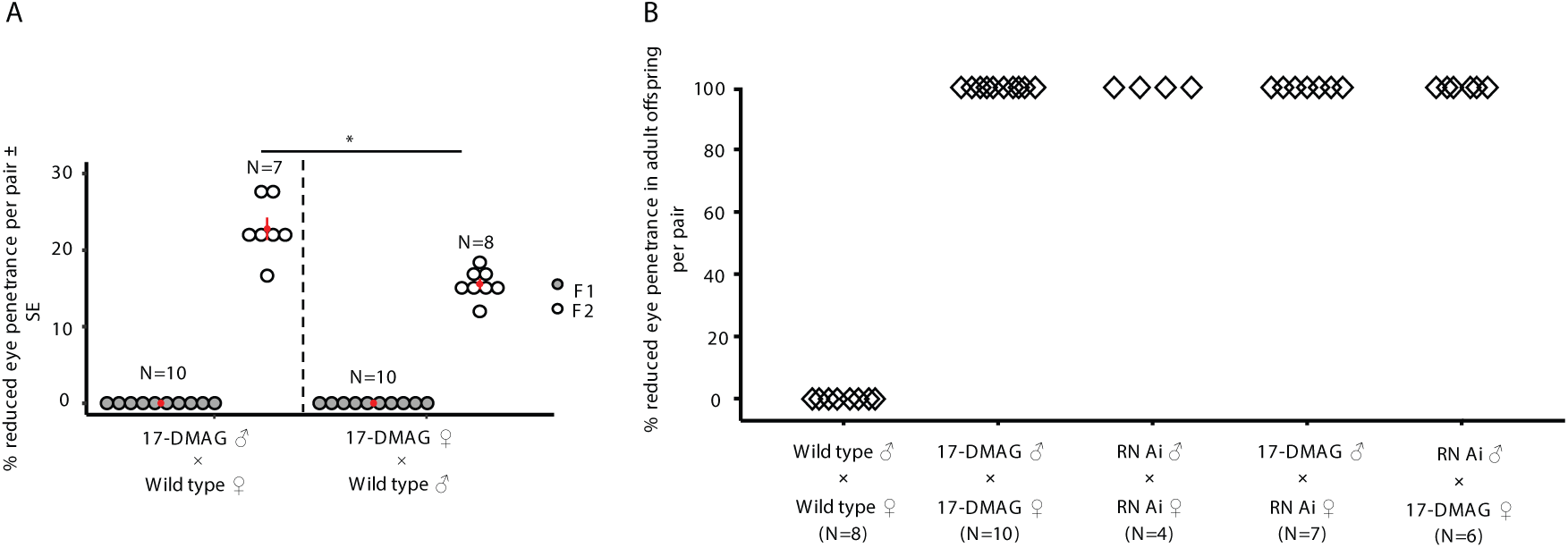
Inheritance of the reduced-eye phenotype. **(A)** Penetrance in F1- and F2-adults after outcrossing the reduced-eye males (♂) and females (♀) with wild type beetles; (GLMM: LRT = 12.57, df = 1, \**p*<0.001). **(B)** Penetrance in adult offspring produced by different crosses between reduced-eye beetles. The reduced-eye beetles were randomly chosen from monomorphic lines established either by 17-DMAG treatment (17-DMAG) or RNAi treatment (RNAi); wild type beetles were randomly taken from the negative control line; each data point represents progeny produced from one family; *N* represents the number of pairs per treatment.

### Fitness of the reduced-eye phenotype

Organisms can respond to environmental changes by expressing adaptive phenotypic variation. An HSP90 buffering role might be one of the specific mechanisms that reveal variation under stress. While most morphological changes controlled by HSP90 are expected to have serious fitness costs, a rare beneficial change might have the potential to evolve free of deleterious fitness effects (Carey et al., 2006). To examine whether the expression and fitness of the reduced-eye phenotype depends on environmental conditions, we used the polymorphic line described above. We exposed unaffected beetles from this line to continuous darkness, continuous light, and mild heat stress, and then compared the proportions of offspring with reduced eyes in the adult stage. There was a significantly higher penetrance of the reduced-eye phenotype under continuous light compared to continuous darkness and heat conditions (Figure 4A).

**Figure 4.**
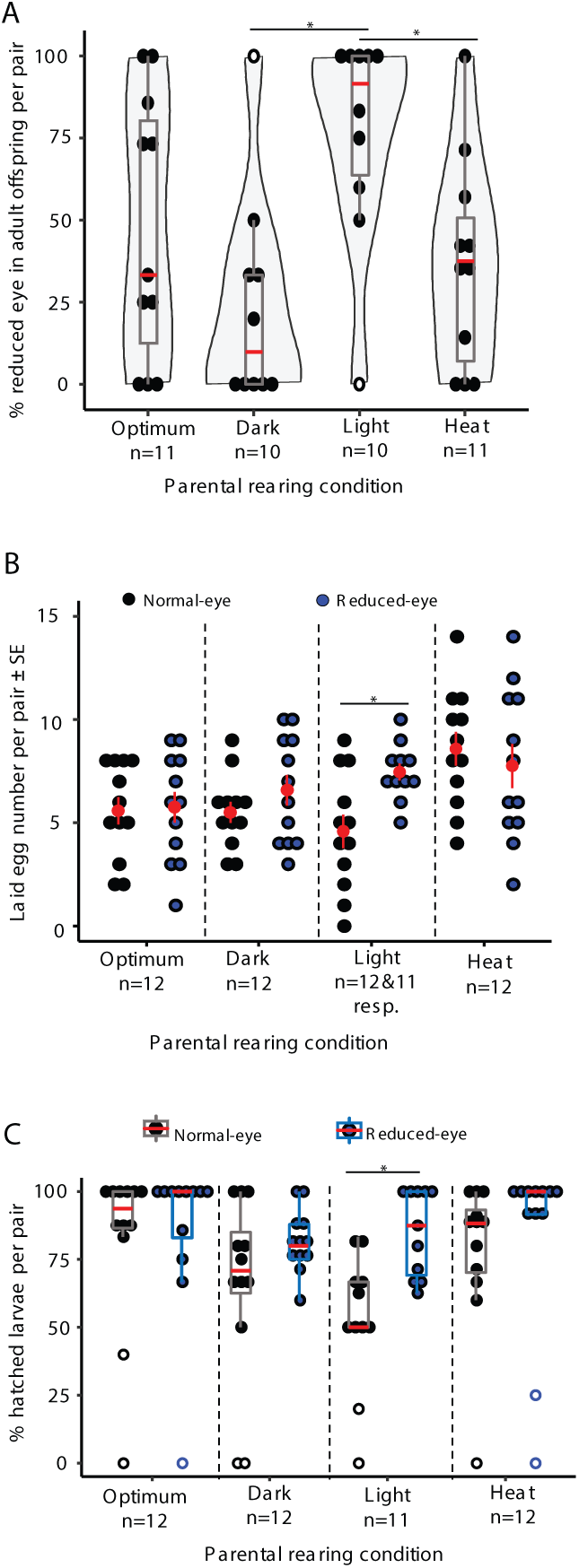
Penetrance and fitness of the reduced-eye phenotype under environmental stress. Beetles were kept for 48 h under optimum conditions (12h/12h dark/light cycle and 30°C), in continuous dark (Dark), continues light (Light) or at 35°C (Heat). (**A**) Penetrance of the reduced-eye phenotype; each data point represents offspring produced from one pair (Kruskal Wallis nonparametric test: Chisq = 10.004, df = 3, *p* = 0.019). (**B**) Numbers of eggs laid by normal and reduced-eye adults in a 24-h egg laying period; mean ± SE in red, (LM: Optimum, F_1,22_ = 0.03, *p* = 0.870; Dark, F_1,22_ = 1.44, *p* = 0.244; Light, F_1,21_ = 9.23, *p* =0.006 and Heat, F_1,22_ = 0.39, *p* = 0.541). (**C**) Proportion of hatched larvae from eggs laid by normal- and reduced-eye adults, (Kruskal Wallis nonparametric test: Optimum, Chisq = 0.23, df = 1, *p =* 0.628; Dark, Chisq = 1.50, df = 1, *p* = 0.221; Light, Chisq = 9.55, df = 1, *p* = 0.002 and Heat, Chisq = 3.13, df = 1, *p* = 0.077). *N* represents the number of pairs; for the boxplots: horizontal lines represent median (red) and inner quartile ranges (25^th^ to 75^th^ percentiles) and vertical lines represent upper and lower whiskers (1.5× interquartile ranges), open dots represent outliers; significant differences indicated by asterisks, *p*<0.05.

We then compared the fitness of beetles with reduced eyes to the unaffected siblings (thereby controlling for genetic background). Under light stress, beetles with reduced eyes laid significantly more eggs than did the normal-eye siblings under the same conditions (Figure 4B), and a higher proportion of larvae hatched from these eggs in the reduced-eye group than in the normal-eye sibling group (Figure 4C).

## Discussion

More than 70 years after Waddington conceived his concept of canalization (Lack et al., 2016; Waddington, 1942), and 20 years after Rutherford and Lindquist’s demonstration of the release of phenotypic diversity upon HSP90 reduction (Rutherford and Lindquist, 1998), the evolutionary relevance of capacitance remains controversial. With the present study, using the important insect model *T. castaneum* (Brown et al., 2009) as a promising system for studying these questions, we demonstrate that a phenotype that appeared after HSP90 impairment had a fitness benefit under specific environmental conditions.

Rutherford and Lindquist’s original idea was that HSP90 serves as an evolutionary capacitor due to its function as a chaperone that acts at the protein level; genetic variants coding for potentially misfolded proteins might remain cryptic, but only as long as HSP90 can fulfill its role as a helper for correct protein folding (Lindquist, 2009). That HSP90 inhibition might also produce new genetic variants through the activation of TEs has been suggested as an alternative (Specchia et al., 2010). The important difference between these two concepts lies in the fact that according to Rutherford and Lindquist, genetic variants that have been proven successful under certain conditions can remain in a population’s gene pool, whereas the vast majority of new genetic variants produced by an HSP90 mutator effect through TE activity are likely to be harmful.

In the present study, the reduced-eye phenotype arose independently, after reduction of HSP90 by two different experimental approaches (RNAi and chemical inhibition), in offspring of individuals stemming from the same starting populations. Furthermore, using crosses, we found evidence that one and the same recessive locus might cause this phenotype. It is rather unlikely that TEs have inserted within the same gene or regulatory region in both experiments. On the other hand, a cryptic reduced-eye gene residing in our starting population could explain why we found the same phenotype independently. Of course, this does not exclude that TEs are relevant for other phenotypes released under HSP90 reduction.

Our crosses are mostly consistent with a single recessive locus. However, penetrance of the reduced-eye phenotype was somewhat reduced when affected females instead of males were used for the crosses (Figure 3A), and it was also slightly influenced by the environment (Figure 4A). Taken together, this indicates that epigenetic processes are involved in the development of the reduced-eye trait, as has previously been suggested for HSP90-mediated processes in other animals (Gore et al., 2018; Sollars et al., 2003; Zabinsky et al., 2019).

Irrespective of the underlying mechanisms, a counter argument against the evolutionary relevance of capacitance has been that most of the phenotypes that were found after HSP90 inhibition likely have deleterious effects on fitness (Carey et al., 2006). However, Queitsch et al. (Queitsch et al., 2002) convincingly argue that some of the phenotypes they found after HSP90 inhibition are not monstrous and might be advantageous under particular conditions, but they did not directly test their fitness. Chen et al. (Chen et al., 2012) demonstrated that inhibiting HSP90 can potentiate fast adaptation to cytotoxic compounds by inducing aneuploidy. Very recently, Koubkova-Yu et al. (Koubkova-Yu et al., 2018) found that by replacing the native HSP90 in *Saccharomyces cerevisiae* cells with the ortholog from hypersaline-tolerant *Yarrowia lipolytica*, the evolved clones showed a wider range of phenotypic variation than cells carrying native HSP90 and some of them exhibited beneficial mutations with a higher fitness improvement.

As discussed by Rohner et al. (Rohner et al., 2013), cryptic variation in eye-size of surface populations of the cavefish *A. mexicanus* released by HSP90 inhibition most likely produces an evolutionary benefit under conditions of darkness, as developing a fully functional, large eye is a costly process (Moran et al., 2015; Niven, 2015). However, fitness was not directly studied in this environment. We thus studied potential fitness benefits of the reduced-eye phenotype in *T. castaneum* and indeed found superior fitness under certain environmental conditions. At first glance, it might appear surprising that fitness benefits manifested under constant light rather than constant darkness. However, as a stored-product pest, *T. castaneum* beetles are photonegative (Misra R. and Englert D., 1985; Park, 1934), and constant light represents a stressful condition that the optical system is not adapted to, thereby presenting a potential benefit to evolving smaller eyes. Interestingly, eye phenotypes were often affected by HSP90 inhibition in several species (Rohner et al., 2013; Rutherford and Lindquist, 1998; Yeyati et al., 2007), raising the possibility that the buffering by HSP90 is limited to specific morphological traits. Further studies to clarify which genes are affected in our and other systems might provide more insights into this question.

In a previous study in *T. castaneum*, we demonstrated that the expressions of *Hsp83* and *Hsp90* were reduced when beetles were kept in an environment that might be perceived as risky, for example, together with wounded conspecifics (Peuss et al., 2015). Together with the present study, a scenario emerges that links evolutionary capacitance and thus evolvability of a population to environmental conditions. Further studies are needed to determine the relevance of this process, in particular under ecological conditions where rapid adaptation based on standing genetic variation is required, such as coevolution, insecticide resistance, or climate change.

## Materials and Methods

### Model organism

Stocks of the red flour beetle, *Tribolium castaneum* (Coleoptera: Tenebrionidae) (*Cro1* strain) were maintained as non-overlapping generations under standard rearing conditions (30°C and 70% relative humidity with a 12 h light/dark cycle) in wheat flour with 5% brewer’s yeast (Milutinović et al., 2013). This beetle strain was collected in Croatia in 2010 and allowed to adapt to laboratory conditions for at least 20 generations before starting any experiments (Milutinović et al., 2013).

### RNAi-mediated *Hsp83* knockdown

*Hsp83* dsRNA was produced to check the release of genetic variation, and *Hsp83-*specific primers were designed (sequences shown in Table supplement 1) on CDS region nucleotides 508-964 (457 bp) of the Tc-Hsp83 gene. The *Hsp83* fragment was cloned into the pZERrOTM-2 vector to add T7 and T7-SP6 promoter sequences, then double-stranded RNA was synthesized using a T7 MEGAscript Kit (Ambion). Parental RNAi was performed by injecting 3-day-old pupae which were randomly chosen from the wild type stock, as previously described (Posnien et al., 2009), with dsRNA at 20 (low) and 100 (high) ng/µl adjusted by NanoPhotometer™ Pearl (Implen, Germany). Before injection, pupae were individualized during their larval stage to avoid cannibalism. As an off-target control, dsRNA specific for asparagine synthetase A (*AsnA*), a specific gene of the bacterium *Escherichia coli* was used, and naïve beetles were used as the negative control. Injected pupae were kept individually until eclosion in 5% yeasted flour under standard rearing conditions.

### Pharmacological inhibition of HSP90

The water-soluble geldanamycin analogue 17-Dimethylaminoethylamino-17-demethoxygeldanamycin, (17-DMAG; InvivoGEN) was used at 10 (low) and 100 (high) µg/ml to inhibit HSP90 function. Flour discs (i.e., thin layers of flour) containing 17-DMAG were made as previously described (Milutinović et al., 2013) with slight modifications. 17-DMAG solution at the desired concentration was mixed well with flour (0.25 g/ml), pipetted into 96-well plates (40 µl/well), and left to dry for 5 hours at 30°C in an oven with air circulation. Larvae (12-16 days old) were randomly chosen from the wild type rearing stocks and allowed to feed on the flour discs and kept in the dark due to the light sensitivity of 17-DMAG. Fresh drug was introduced every other day, 6 times or until pupation. Flour discs without 17-DMAG were used as a negative control. The eclosed virgin adults were fed on normal loose flour under standard rearing conditions.

### RT-qPCR

To confirm the potential effect of RNAi and 17-DMAG, we used quantitative real-time polymerase chain reaction (RT-qPCR) analysis according to a previously described protocol (Peuss et al., 2015); RT-qPCR primers are shown in Table supplement 1. For RNAi, *Hsp83* expression was measured in the eclosed adults, four biological replicates of pools of 5 individuals, taken 7 days after their pupal injection. For 17-DMAG, the expression of *Hsp68a* was used as a molecular marker for HSP90 inhibition after 17-DMAG treatment (Kudryavtsev et al., 2017; Zhou et al., 2013). Hsp68a expression was measured in individuals collected at different time points of feeding on flour discs containing 17-DMAG at 100 µg/ml (on days 2, 4, 7, 9, and 11, and on day 18, one week after the last feeding time. Four biological replicates of pools of 8 individuals were used for each time point. Gene expression in all experiments was computed relative to the expression of the housekeeping genes *ribosomal protein L13a* (*RpL13a*) and *ribosomal protein 49* (*rp49*).

### Documentation of morphological variation in untreated, subsequent generations after parental HSP90 impairment

For parental *Hsp83-*RNAi treatment, 5-day-old, eclosed beetles were crossed as single pairs. 20-30 single pairs per treatment were allowed to lay eggs for 3 days. Untreated, 10-day-old F1-larvae were individualized to avoid cannibalism, morphologically inspected under the binocular at 15 days old, and then re-individualized until eclosion. Three weeks later, they were re-examined as adults for phenotypic variation. The larvae that showed abnormal phenotypes were kept separately to follow up their development as adults. Beetles (7 days old) that eclosed after larval 17-DMAG treatment were crossed as single pairs (6 to 13 families per treatment) and allowed to lay eggs for 7 days; the untreated F1-adults were then morphologically inspected (body size, color, head, thorax, abdomen, antennae, eyes, mouthparts, legs, and urgomophi). Images of interesting traits were captured, after light anesthesia with CO_2_, by a Canon digital camera attached to a Zeiss Axioskop compound microscope.

### Quantitative trait measurement

Some adult beetles exhibited a reduced-eye phenotype. To measure the eye size, these beetles were CO_2_-anesthetized, kept at a fixed position on their ventral side, and photographed with a Canon digital camera attached to a Zeiss Axioskop compound microscope. ImageJ was used to measure the eye area of both left and right sides. The body length was measured dorsally and ventrally from the anterior end of the head to the posterior end of the abdomen, and the mean of both sides represented the body length.

### Production of lines with specific traits

For both RNAi and 17-DMAG treatments, the untreated, affected F1-adult beetles that showed morphological abnormalities were selected and mated as single pairs (for RNAi: *N* = 11-19 F1-pairs, most exhibited leg phenotypes; for 17-DMAG: *N* = 18-29 F1-pairs, some exhibited the reduced-eye phenotype) for each treatment group to obtain F2 generations. F2 adults were morphologically examined, focusing on the trait of interest in addition to any other new traits. F2 individuals that exhibited specific traits were mated in groups rather than single pairs, to increase the population size and create large numbers of individuals, enabling us to study the inheritance of certain morphological variations over generations. To establish a ‘polymorphic line’ from the RNAi treatment, beetles that showed interesting traits in the F2 generation were pooled with their normal siblings, without any artificial selection, to produce an F3 generation, and the trait was checked until the 7^th^ generation.

For the 17-DMAG treatment group, we established a ‘monomorphic line’ by crossing only the reduced-eye F2 beetles. We also established a ‘polymorphic line’ by crossing their normal siblings. The lines were bred under normal conditions without any artificial selection, and the trait was checked every generation up to F7.

### Penetrance and fitness of the reduced-eye phenotype under environmental stress

We studied the penetrance and fitness of reduced-eye beetles under normal and stress conditions and compared them to their normal eye siblings, which were produced from the same grandparents and therefore had the same genetic background. Beetles were randomly selected from the F5 generation of the polymorphic line originally established by 17-DMAG treatment. Virgin adult beetles were kept under either standard- or stress-conditions, including continuous dark, continues light, and heat stress at 35°C and 70% relative humidity. Beetles were stressed for 24 hours, then crossed as single pairs (*N* = 11-12) and allowed to lay eggs for 24 hours under the same stress conditions (Hawk et al., 1974). Oviposition and hatchability were used as fitness measures; egg number was checked 1 day after mating and the number of hatched larvae was investigated 10 days later. Penetrance of the reduced-eye phenotype was documented in adult offspring 34 days after egg laying.

### Statistical analyses

#### Morphological variation

For RNAi and 17-DMAG experiments (Figures 1A, 1C, 2A and Figure 1-figure supplement 2B), the significant differences in abnormal phenotypes for either larvae or adults were tested using a generalized linear model (GLM) with a binomial error distribution in JMP (JMP^®^, version 11, SAS Institute). The state of animals was used as a response variable, each individual was given a value of 1 (affected) or 0 (not affected), and the pairing treatment groups were used as an independent variable. A post hoc contrast test was used to test against the control groups.

In the experiment of outcrossing with wildtype beetles (Figure 3A), the reduced eye penetrance in the F2 generation was compared in the two different pairing groups (after exclusion of two outliers above 95th percentile which were considered to be random noise in all distributions, and not considered for the analysis) using a GLMM model in R Version 3.2.1 as follows: Model: Number of reduced-eye F2-beetles contrasted with normal-eye F2-beetle number _per family_ ∼ Treatment _Reduced-eye ♂× WT ♀, Reduced-eye ♀× WT♂_/ Family number _random_).

To statistically analyze the penetrance of the reduced-eye phenotype in offspring under normal and stress conditions (Figure 4A), the significant differences were checked only in the offspring produced from normal eye siblings. We excluded from the analysis the reduced-eye group, as they exhibited eye penetrance of 100% under all rearing conditions in all families. Non-parametric comparisons were performed using the Kruskal–Wallis (Rank Sums) test in R Version 3.2.1 using the ‘agricolae’ package, as the residuals of GLMs were not normally distributed (i.e., exhibited overdispersion; Model: Reduced eye percent_per family_ ∼ Rearing condition). A post-hoc pairwise test was performed in the ‘multcompView’ package to determine which levels of the fixed variable differed from one another.

#### Trait analysis

To evaluate measurement error, eye size of all individuals was measured twice to test for repeatability (*R*) of measurements (Arnqvist and Mårtensson, 1998). As our measurements were highly repeatable (*R* > 0.9), the eye size of each individual (Figure 2B and Figure 1-figure supplement 2A) was represented by the mean of the two measures and significant differences in eye size (expressed as means ± SEM) among treatment groups. The relative eye size in Figure 1-figure supplement 2A was statistically analyzed using unequal variances analysis, two-tailed F-test, Levene’s test, and Bartlett’s test using JMP (JMP^®^, version 11, SAS Institute), where the response variable was the relative eye size (calculated by dividing the eye size in mm^2^ by the body length in mm) in each individual and the independent variable was the parental treatment group.

#### Fitness measures

The oviposition and hatching rate of reduced-eye beetles that were maintained under different rearing conditions were compared to those of their normal-eye siblings. We measured changes in the number of laid eggs (Figure 4B) using a linear model in R, ‘MASS’ package, followed by Tukey’s HSD test as a pairwise test. The response variable was the egg number laid by each family within treatments and the independent variable was the beetle group. We ran a separate model for each rearing condition according to the following: Model: Egg number per family ∼ Beetle groups (reduced & normal-eye groups) _optimum, dark, light or heat_.

The numbers of hatched larvae (Figure 4C) were statistically compared using a nonparametric Kruskal-Wallis test (Rank Sums) in R using the ‘agricolae’ package, as the GLM residuals were overdispersed. The percent hatchability for each family was used as a response variable and the independent variable was the beetle group under a certain rearing condition; outliers were included in the statistical analysis; separate models were run for each rearing condition: (Model: Hatchability percent _per family_ ∼ Beetle groups (reduced- & normal-eye groups) _optimum, dark, light or heat_). Post hoc analysis of significant differences was conducted using the Dunn test. Individuals that did not lay any eggs were excluded from this analysis.

## Acknowledgements

We would like to thank Jürgen Schmitz and Gennady Churakov (ZMBE Münster) as well as Robert Peuß (Stowers Institute for Medical Research, Kansas City, USA) for helpful suggestions and discussions, Lukas Schrader (Institute for Evolution and Biodiversity, University of Münster, Münster, Germany) for commenting on our manuscript, and Barbara Hasert for technical assistance. Funding: This research was partly funded by the Deutsche Forschungsgemeinschaft (DFG, German Research Foundation) – Projektnummer 316099922 – TRR 212 and a doctoral stipend from the DAAD to RA.

## Author contributions

RA and JK designed the study. RA performed the experiments. RA and JK wrote the manuscript.

## Data availability

Raw data are attached to our article.

## List of Figure supplement legends

**Figure 1-figure supplement 1.**
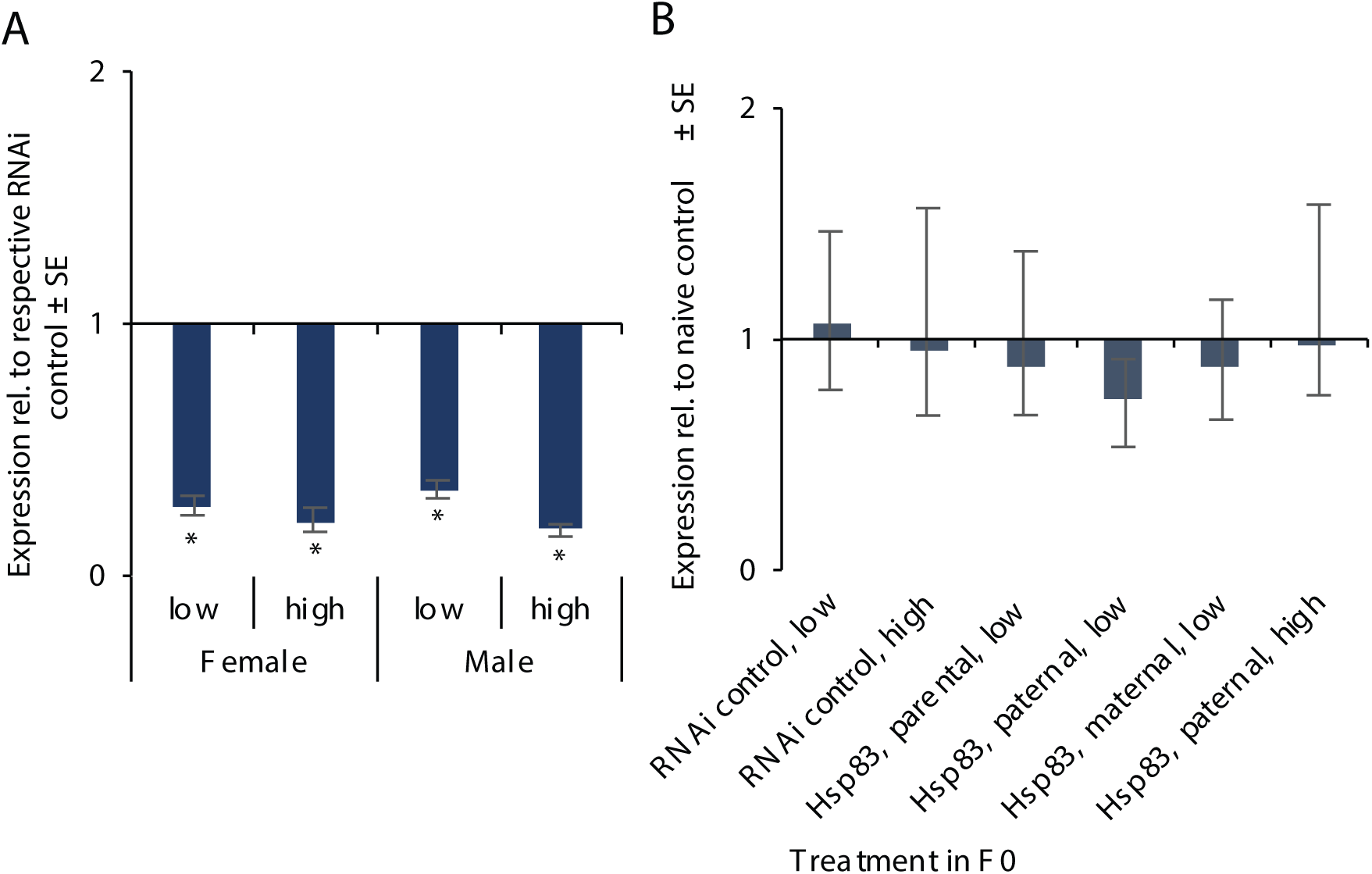
Relative expression of the Hsp83 gene using RT-qPCR. **(A)** Expression in eclosed adults (females and males) after *Hsp83* knockdown by RNAi. Pupae were injected with 20 (low) or 100 (high) ng/μl *Hsp83* dsRNA; *n* = 4 biological replicates, each with a pool of five beetles. **(B)** Expression in untreated F1 larvae (10 days old) produced from *Hsp83* knockdown parents (F0) relative to those produced from untreated parents; *n* = 4 biological replicates, each with a pool of ten larvae. The expression is relative to t hose in the housekeeping genes *RpL13a* and *Rp49*; data are means ± SEM calculated by REST 2009; * *p*<0.005.

**Figure 1-figure supplement 2.**
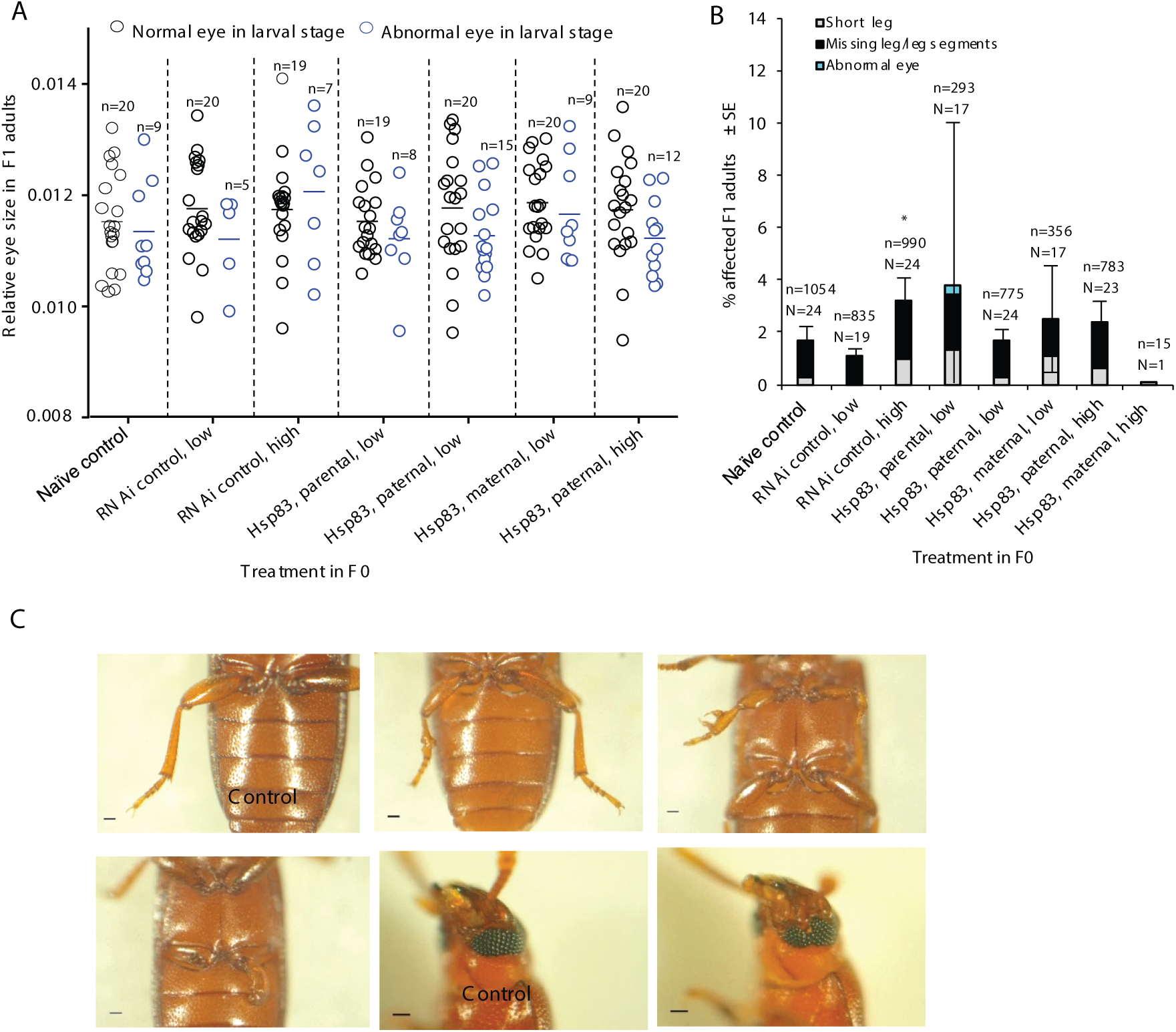
Morphological variations in F1 offspring of HSP90-inhibited parents (RNAi-mediated knockdown of *Hsp83* with 20 (low) and 100 (high) ng/μl dsRNA). **(A)** Quantification of eye size relative to body length in F1 adults that exhibited either normal-eye (black circles) or abnormal-eye (blue circles) phenotypes during their larval stage; each data point represents the relative eye size for one individual and *n* shows the total number of individuals. There were no significant differences in eye size among treatments or even within treatments (Unequal variances analysis, two-sided F test: F = 1.085, df = 13, *P* = 0.391; Bartlett’s test, *P* = 0.686; Levene’s test, *P* = 0.534); horizontal lines represent means. **(B)** Proportion of affected adults (total number of *n* screened individuals produced from *N* pairs per treatment shown above each bar; * *p*<0.05, relative to naïve control; see details in Table S1). **(C)** example phenotypes: short and missing legs and abnormal eyes; scale bars: 100 µm.

**Figure 1-table supplement 1.** Statistical analyses of morphological variations in subsequent generations produced after HSP90 impairment using RNAi. RNAi-mediated knockdown of *Hsp83* was carried out using dsRNA at 20 (low) and 100 (high) ng/μl. The generalized linear model (GLM) was applied using a binomial error distribution in which each individual was given a value of 1 (affected) or 0 (not affected). A post hoc contrast test was used to compare the treatment groups against the naïve control group; *p*-values less than 0.05 are in bold.

| Tested groups | untreated F1-larvae |  |  | untreated F1-adults |  |  | untreated F2-adults |  |  |
| --- | --- | --- | --- | --- | --- | --- | --- | --- | --- |
| | df | L-R $\chi^2$ | *p* | df | L-R $\chi^2$ | *p* | df | L-R $\chi^2$ | *p* |
| RNAi control, parental, low | 1 | 1.373 | 0.241 | 1 | 1.346 | 0.246 | 1 | 1.833 | 0.176 |
| RNAi control, parental, high | 1 | 2.482 | 0.115 | 1 | 5.021 | **0.025** | 1 | 2.282 | 0.131 |
| *Hsp83*, parental, low | 1 | 25.750 | **<0.0001** | 1 | 2.510 | 0.088 | 1 | 6.538 | **0.011** |
| *Hsp83*, paternal, low | 1 | 41.334 | **<0.0001** | 1 | 0.009 | 0.924 | 1 | 10.968 | **0.001** |
| *Hsp83*, maternal, low | 1 | 1.282 | 0.257 | 1 | 0.79 | 0.374 | 1 | 7.571 | **0.006** |
| *Hsp83*, paternal, high | 1 | 36.067 | **<0.0001** | 1 | 1.009 | 0.315 | 1 | 43.954 | **<0.0001** |
| *Hsp83*, maternal, high | 1 | 0.119 | 0.73 | 1 | 1.146 | 0.284 | 1 | 0 | 1 |
| Among groups | 7 | 85.008 | **<0.0001** | 7 | 15.509 | **0.03** | 7 | 77.608 | **<0.0001** |

**Figure 2-figure supplement 1.**
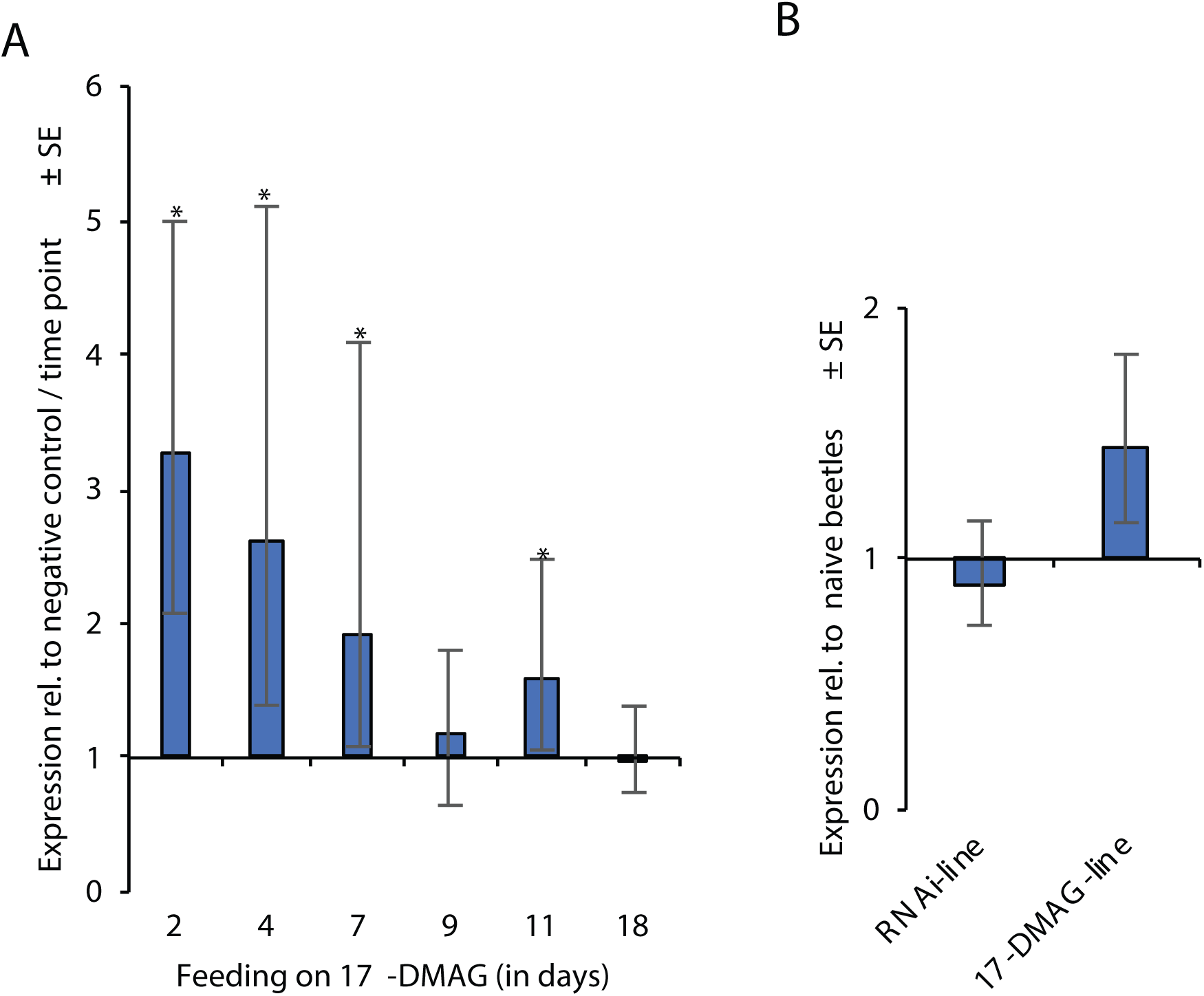
Relative expression of heat shock protein genes using RT-qPCR. **(A)** Expression of the Hsp68a gene as a molecular marker for the effect of 17-DMAG. Expression was measured in larvae fed on flour discs containing 100 μg/ml of 17-DMAG on days 2, 4, 7, 9, and 11 and in eclosed adults one week after the last feeding time (day 18); *n* = 4 biological replicates of pools of 8 individuals; expression at each time point is compared to the negative control (flour discs without 17-DMAG) at the respective time point. **(B)** *Hsp83* expression in reduced-eye beetles of lines established after HSP90 impairment using either RNAi or 17-DMAG in the parental generation; *n* = 3 biological replicates of pools of five beetles. The expression is relative to t hose in the housekeeping genes *RpL13a* and *Rp49*; data are means ± SEM calculated by REST 2009; * *p*<0.005.

**Figure 2-figure supplement 2.**
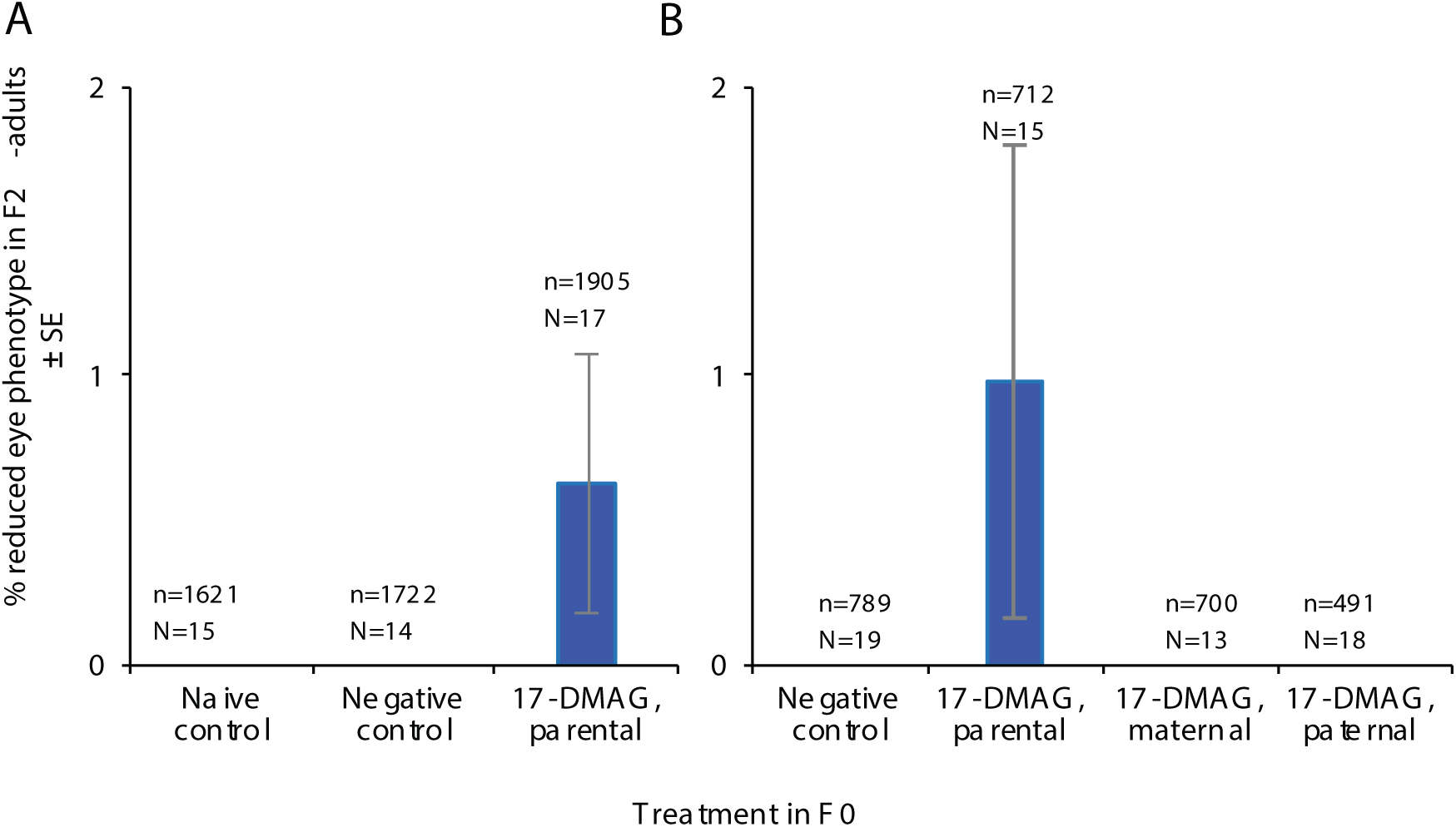
Proportion of reduced-eye trait in F2 adults measured in two independent experiments (A) and (B). F2 beetles were produced after HSP90 inhibition in the parental generation (F0) by 17-DMAG at 100 μg/ml. Total number of *n* screened individuals produced from *N* pairs per treatment shown above each bar. **(A)** Twelve of 1905 F2-beetles exhibited the reduced-eye phenotype; all 12 of these beetles resulted from two F1-pairs (6.8%; 6/88) and (3.8%; 6/156). **(B)** Seven of 712 F2-beetles exhibited the reduced-eye phenotype; all seven beetles were produced from one F1-pair (12.3%; 7/57).

**Figure 2-table supplement 1.** Statistical analyses of morphological variations in F1-adults produced after HSP90 impairment using 17-DMAG. HSP90 chemical inhibition was mediated by 17-DMAG at 10 (low) and 100 (high) μg/ml. The generalized linear model (GLM) was applied using a binomial error distribution in which each individual was given a value of 1 (affected) or 0 (not affected). A post hoc contrast test was used to compare the treatment groups against the naïve control group; *p*-values less than 0.05 are in bold.

| Tested groups | untreated F1-adults |  |  |
| --- | --- | --- | --- |
| | df | $L\text{-}R\ \chi^2$ | $p$ |
| Negative control | 1 | 2.966 | 0.085 |
| 17-DMAG, low | 1 | 4.517 | **0.034** |
| 17-DMAG, high | 1 | 29.887 | **<0.0001** |
| Among groups | 3 | 32.742 | **<0.0001** |

**Table supplement 1.** Primers used in this study

